# Scent dog identification of SARS-CoV-2 infections, similar across different body fluids

**DOI:** 10.1101/2021.03.05.434038

**Authors:** Paula Jendrny, Friederike Twele, Sebastian Meller, Claudia Schulz, Maren von Köckritz-Blickwede, Ab Osterhaus, Hans Ebbers, Janek Ebbers, Veronika Pilchová, Isabell Pink, Tobias Welte, Michael Peter Manns, Anahita Fathi, Marylyn Martina Addo, Christiane Ernst, Wencke Schäfer, Michael Engels, Anja Petrov, Katharina Marquart, Ulrich Schotte, Esther Schalke, Holger Andreas Volk

## Abstract

**Background:** The main strategy to contain the current SARS-CoV-2 pandemic remains to implement a comprehensive testing, tracing and quarantining strategy until vaccination of the population is adequate.

**Methods:** Ten dogs were trained to detect SARS-CoV-2 infections in beta-propiolactone inactivated saliva samples. The subsequent cognitive transfer performance for the recognition of non-inactivated samples were tested on saliva, urine, and sweat in a randomised, double-blind controlled study.

**Results:** Dogs were tested on a total of 5242 randomised sample presentations. Dogs detected non-inactivated saliva samples with a diagnostic sensitivity of 84% and specificity of 95%. In a subsequent experiment to compare the scent recognition between the three non-inactivated body fluids, diagnostic sensitivity and specificity were 95% and 98% for urine, 91% and 94% for sweat, 82%, and 96% for saliva respectively.

**Conclusions:** The scent cognitive transfer performance between inactivated and non-inactivated samples as well as between different sample materials indicates that global, specific SARS-CoV-2-associated volatile compounds are released across different body secretions, independently from the patient’s symptoms.

**Funding:** The project was funded as a special research project of the German Armed Forces. The funding source DZIF-Fasttrack 1.921 provided us with means for biosampling.

## Introduction

### Current situation

The recently emerged respiratory disease coronavirus disease 2019 (COVID-19) broke out in Wuhan, Hubei Province of the People’s Republic of China, for the first time in December 2019 and was declared a global health emergency by the World Health Organization at the end of January 2020^1,2^. It rapidly developed into a global pandemic within just a few months. The pandemic has led to enormous restrictions and sanctions affecting public as well as private life. The severe acute respiratory syndrome coronavirus 2 (SARS-CoV-2), which causes COVID-19, infects the upper respiratory tract and in more serious cases may also cause severe pneumonia and acute respiratory distress syndrome. The clinical presentation of SARS-CoV-2 infection is heterogeneous, ranging from asymptomatic infection to typical symptoms such as fever, cough, fatigue, ageusia and anosmia, but may also present atypically and lead to multiorgan dysfunction and death^1,3,4^. Containing this global pandemic requires a high rate of testing, as an effective tool to contain viral spread. Viral loads can be detected by reverse transcription polymerase chain reaction (RT-PCR) assays and with slightly less sensitive and usually more rapid antigen detection tests in nasal or pharyngeal swabs^2,4^ and saliva^5,6,7^ with a peak at days three to ten after infection. The peak of infectiousness is around symptom onset^8^. It remains unclear if sweat or urine are also sources of virus transmission^9,10^.

### Odour detection

Different infectious diseases may cause specific odours by emanating volatile organic compounds (VOCs). These are metabolic products, primarily produced by cell metabolism and released through breath, saliva, sweat, urine, faeces, skin emanations and blood^11^. The VOC-pattern reflects different metabolic states of an organism, so it could be used for medical diagnosis by odour detection and disease outbreak containment^12^.

Canines are renowned for their extraordinary olfactory sense, being deployed as a reliable tool for real-time, mobile detection of, e.g., explosives, drugs and may identify certain pathogen- and disease-specific VOCs produced by infected body cells. The limit of detection for canines is at concentrations of one part per trillion, which is three orders of magnitude more sensitive than currently available instruments^12^. Consequently various studies have shown dogs’ abilities to detect with high rates of sensitivity and specificity^13^ infectious and non-infectious diseases and conditions, such as different types of cancer^14^, malaria^15^, bacterial infections caused by e.g. *Clostridium difficile* or mastitis causing pathogens^16,17^, hypoglycaemia in diabetics^18^, and virus infections in cell cultures^12,19^. In addition, several research groups worldwide currently train and deploy SARS-CoV-2 detection dogs^20,21^. In a pilot study, our group showed that dogs were able to detect inactivated saliva samples from COVID-19 patients with a sensitivity of 83% and specificity of 96%^22^, which has been confirmed by other groups training dogs to detect either respiratory secretions or sweat samples from COVID-19 patients^20,21^. Despite these preliminary promising results, it remains to be shown whether dogs detect VOCs which are biofluid-specific or alternatively there is a more general change in odour of COVID-19 patients. To test the latter hypothesis, the current study used the same training set-up with inactivated saliva samples as the former study^22^ and investigated whether dogs could transfer their smell recognition to non-inactivated saliva, urine or sweat samples from SARS-CoV-2 infected patients. Scent detection dogs could be a reasonable option for a first line screening method in public facilities such as airports or during major events as well as in retirement homes or medical institutions that would be real-time, effective, economical, effortless and non-invasive.

## Methods

### Samples - target scent, negative controls and distractors

To acquire saliva samples, individuals had to salivate about 1-3 ml through a straw into sample tubes. For the training phase, saliva samples from twelve subjects (hospitalised and non-hospitalised SARS-CoV-2 infected individuals) suffering from asymptomatic to severe COVID-19 symptoms were inactivated with beta-propiolactone (BPL) according to the protocol described in Jendrny et al. 2020 to provide safe training conditions for dogs and handlers. To generate sweat samples, the test persons had to wipe their crook of the arm with a cotton pad. Urine samples were collected from the test persons by urinating into a cup and transfer of 5 ml into a sample tube. After acquisition, all of the samples were deep-frozen at - 80°C in the laboratory until usage. Samples from ninety-three participants were used in the study (**suppl. table 1**). The SARS-CoV-2 status of each collected sample was determined by the RT-PCR SARS-CoV-2-IP4 assay from Institut Pasteur including an internal control system and protocol^22,23^.

In contrast to our first study^24^, which only included hospitalised COVID-19 patients suffering from severe courses of disease, we now additionally included non-hospitalised asymptomatic individuals as well as individuals with mild clinical signs. Inclusion criteria were either the diagnosis of infection by positive SARS-CoV-2 RT-PCR of nasopharyngeal swabs (positive samples), negative SARS-CoV-2 RT-PCR result and healthy condition (negative control samples) or negative SARS-CoV-2 RT-PCR result and symptoms of other respiratory disease (distractor samples). Written consent from all participants were collected before sample collection. The local Ethics Committees of *Hannover Medical School* (MHH) and the Hamburg Medical Association (for the *University Medical-Center Hamburg-Eppendorf* (UKE)) approved the study (ethic consent number 9042_BO_K_2020 and PV7298, respectively).

To ensure safety for presentation of non-inactivated samples, glasses specially designed for scent dog training (Training Aid Delivery Device (*TADD*), Sci-K9, USA) containing an odour-permeable but hydro- and oleophobic fluoropolymer membrane were used. A 1 x 1 x 0.5 cm cotton pad soaked with 100 µl of fluid sample material or a snippet of the cotton pad used for sweat sampling was placed at the bottom of the *TADD*-glass and the glass was safely sealed in the laboratory under biosafety level 3 laboratory conditions.

### Dogs

All ten dogs were German armed forces’ service dogs with a history of either protection work, explosives detection or no previous training except for obedience (**suppl. table 2)**. Involved dog breeds were Malinois (n=5), Labrador Retriever (n=3), German Shepherd (n=1) and a Dutch shepherd crossbreed dog (n=1) with ages ranging between one and nine years (median age= 3.7 years). Six female and four male dogs were included.

### Testing device

For the detection training and testing, a device called ‘Detection Dog Training System’ (DDTS, Kynoscience UG, Hörstel, Germany) was utilised, which provided automated and randomised sample presentations for the dogs as well as automatic rewards as described previously^24^. The recorded results were verified by manual video analysis.

### Training procedure

The training procedure was exclusively based on positive reinforcement. Dogs were familiarised to the DDTS for six days using a replacement odour, followed by specific training for 8 days to condition them for the scent of SARS-CoV-2 infections in twelve inactivated positive saliva samples and negative control samples from healthy individuals, respectively. The final study was conducted on four days (four hours a day) and included non-inactivated saliva samples as well as urine and sweat samples. All of the samples used in the final study had not been presented to the dogs before.

### Study design of the double-blinded study

The study was conducted in compliance with safety and hygiene regulations according to the recommendations of the Robert Koch Institute (Berlin, Germany), and approved by local authorities (regional health department and state inspectorate’s office; Hannover, Germany). All samples were handled by the same person with personal protective equipment including powder-free nitrile gloves to prevent odour contamination which may irritate the dogs. In the first session non-inactivated saliva samples were used to assess whether dogs were able to transfer their trained sniffing performance from inactivated to non-inactivated saliva samples. In the following sessions, the detection performances for non-inactivated sweat, urine, and, again, saliva samples were evaluated. There were four possibilities for the dogs to respond to the presented odours:

1. True positive (TP): the dog correctly indicates a SARS-CoV-2 positive sample
2. False positive (FP): the dog incorrectly indicates a negative control or distractor
3. True negative (TN): the dog sniffs shortly at a negative sample but correctly does not indicate it
4. False negative (FN): the dog sniffs shortly at a positive sample but does not indicate it

A detection trial was considered accomplished if the dog left his snout in the target scent presenting hole for ≥ 2 sec, initiating automatically the next trial. In each trial, the device’s software randomly assigned the target scent’s position between the seven different positions without the dog or its handler knowing which hole was next positive. The results were recorded electronically for subsequent analysis and verified by manual time-stamped video analysis. The standard temperature in the dog training laboratory was controlled at 24 ± 1°C. Although the samples were presented to the dogs in safe specimen vessels (*TADD*-glasses), the detection experiments with infectious material were performed in a biosafety level 2 laboratory to prevent any risk of infection. After leaving the test room, the canines were washed with 4% chlorhexidine shampoo with at least ten min contact time to prevent any potential environmental contamination and virus spread. The equipment was disinfected after each test day with suitable disinfectant wipes soaked in limited virucidal disinfectant solution. In addition, swab samples of the dogs’ noses and from the outside of *TADD*-membranes were taken after each day of testing and examined with RT-PCR-assays at the Central Institute of the Bundeswehr Medical Service or Research Center for Emerging Infections and Zoonoses to exclude contamination and replication with infectious viral particles in the dogs’ noses or an escape of virus-containing material from the vessel (**suppl. table 3**).

### Analysis of sensitivity and specificity

The diagnostic sensitivity as well as diagnostic specificity, positive predictive values (PPV), and negative predictive values (NPV) were calculated according to Trevethan^25^. 95% confidence intervals (CIs) for sensitivity and specificity were calculated with the Wilson Score Method^26^. In addition, medians of sensitivity, specificity, PPV, NPV, and accuracy with corresponding 95% CIs of median were calculated. All calculations were done with the Prism 9 software from GraphPad (La Jolla, CA, USA).

## Results

When non-inactivated saliva samples were presented to the dogs after training with inactivated saliva samples, dogs were able to discriminate between samples of infected (RT-PCR positive), non-infected (RT-PCR negative) individuals and distractor samples (RT-PCR negative but respiratory symptoms) with a diagnostic sensitivity of 84% (95% CI: 62.5–94.44%) and specificity of 95% (95% CI: 93.4–96%). During the following detection sessions, when the device was equipped with non-inactivated samples with the same body fluid (saliva, sweat or urine), the corresponding values for diagnostic sensitivity and specificity for saliva samples were 82% (95% CI: 64.29–95.24%) and 96% (95% CI: 94.95–98.9%), for sweat samples 91% (95% CI: 71.43–100%) and 94% (95% CI: 90.91–97.78%), and for urine samples 95% (95% CI: 66.67–100%) and 98% (95% CI: 94.87–100%) respectively (**Fig. 1, suppl. table 4**).

**Figure 1.**
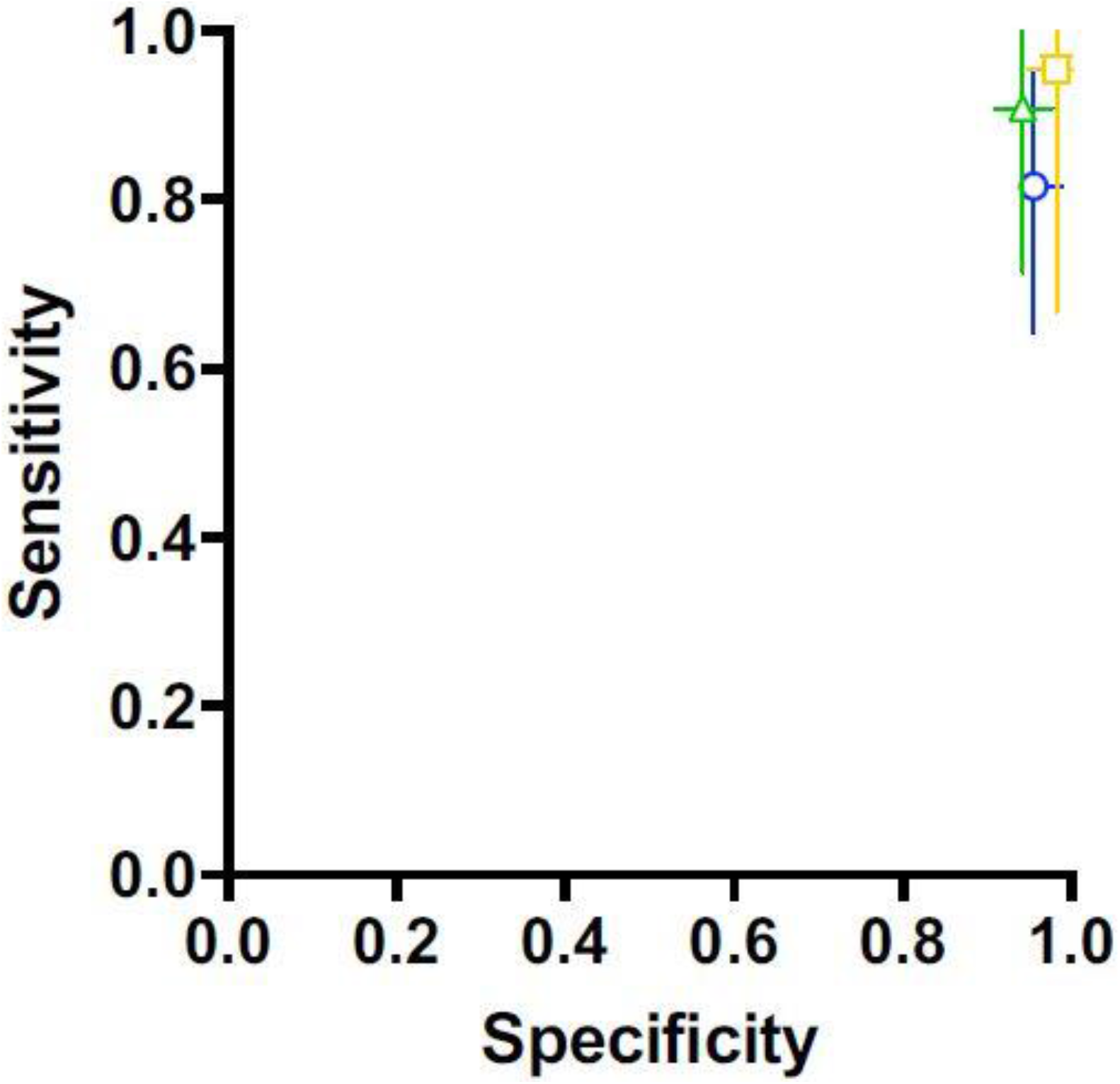
Median diagnostic specificity and sensitivity for all dogs for non-inactivated sweat (green triangle), urine (yellow square), and saliva (blue circle) samples, respectively. The 95% confidence intervals of the medians for specificity and sensitivity are shown with horizontal and vertical bars, respectively.

During the presentation of 5308 randomised sample presentations, the overall success rate was 92% with 723 correct indications of positive, 4140 correct rejections of negative or distractors, 214 incorrect indications of negative and incorrect rejections of 231 positive sample presentations **(Suppl. table 5)**. The RT-PCR results of the sample material from participants with a diagnosed SARS-CoV-2 infection via nasopharyngeal swab and RT-PCR were only positive in twelve cases. Nasopharyngeal swabs from each dog, as well as from the outside of the membranes taken after each day of testing were all negative.

## Discussion

Fast, rapid, affordable and accurate identification of SARS-CoV-2 infected individuals remains pivotal not only for limiting the spread of the current pandemic, but also for providing a tool to limit the impact on public health and the ecomony. Data from the current scent dog detection study confirm our former pilot study (sensitivity 84% versus 83% and specifity 95% versus 96%, respectively). In the current study, dogs were after only eight days of training not only able to immediately transfer their scent detection abilities from inactivated to non-inactivated saliva samples, but also to sweat and urine, with urine having the highest sensitivity of 95% and specificity of 98%. These results suggest a general, non-cell specific, robust VOC-pattern generation in SARS-CoV-2 infected individuals and provide further evidence that detection dogs could provide a reliable screening method providing immediate results.

In the former pilot study from our group^24^, only BPL-inactivated samples of COVID-19 patients and controls were used. The first step in the current trial was therefore to evaluate if dogs can transfer scent recognition to non-inactivated saliva samples, even when trained only with inactivated samples. The inactivation process with BPL did not impair the SARS-CoV-2-associated scent of the samples, as dogs were able to discriminate with a similar accuracy between inactivated and non-inactivated saliva samples from SARS-CoV-2 infected individuals and controls. This has huge implications for the training of dogs, as the health and safety measures other groups had to follow when using non-inactivated samples can be overcome by using BPL-inactivation. Data from the current study indicate that dogs can familiarise to a training device and be safely trained within little more than a week by using inactivated saliva samples from SARS-CoV-2 positive individuals and controls and become reliable SARS-CoV-2 detection dogs for untreated samples. Furthermore, the safety of working with the TADD-glasses was also confirmed by negative PCR results of the samples attained (canine nasopharynx and outer TADD-glas-membrane).

In a second step, untreated (non-inactivated) saliva, sweat and urine samples were presented to the dogs separately to evaluate if they can transfer scent recognition from saliva samples to other untreated body fluids. The detection rate for this experiment was also high, especially considering the dogs having not been trained with sweat or urine samples before. In order to eliminate the risk of recognizing an individual odour from a specific subject, samples used were different for each session.

The sample material of the individuals with positive SARS-CoV-2 status (nasopharyngeal swab tested positive via RT-PCR) was predominantly RT-PCR negative which means that dogs are able to detect the changes in metabolism of non-infectious secretions of SARS-CoV-2 infected individuals. This could explain some of the anecdotal reports from the scent detection work at Helsinki airport that dogs were able to detected asymptomatic SARS-CoV-2 infected individuals prior of them shedding virus.

The fact that dogs were able to discriminate successfully between positive, negative samples and distractors represents evidence of a successful discrimination process, whereas the detection ability across three bodyfluids from 93 different individuals indicates a successful generalisation process. Several research groups that currently also train SARS-CoV-2 sniffer dogs achieved good results, which support this work and consolidate the reliability of the canines’ olfactory sense for medical purposes. Grandjean et al. (2020) trained six dogs in one to three weeks using sweat samples and achieved success rates between 76 to 100%^20^. In addition to their work, where only sweat samples from hospitalised patients were used, the current study suggests that also asymptomatic SARS-CoV-2 infected individuals can be detected by the dogs. Our dogs were able to identify different COVID-19 disease phenotypes and phases of disease expression (sore throat, cough, cold, headache and aching limbs, fever, loss of smell and taste and/or severe pneumonia). Another scent dog detection study conducted by Vesga et al. (2020) achieved promising results (95.5% average sensitivity and 99.6% specificity) and also planned real-life experiments^21^. These studies support the evidence of canines offering a reliable screening method for SARS-CoV-2 infections. Future studies are important to address some remaining limitations such as the low number of distractor samples with specified pathogens (differentiation to other lung diseases or pathogens such as infections with other seasonal respiratory viruses, like influenza viruses, rhinoviruses, respiratory syncytial virus, human metapneumovirus, adenovirus, and coronaviruses other than SARS-CoV-2). This was however not within the scope of the current study. The laboratory identification of the specific VOC pattern is still in its infancy, but some current studies under review and also a peer-reviewed one showed SARS-CoV-2 specific biomarkers in breath samples detectable by gas chromatography-ion mobility spectrometry^27,28^, which also support our hypothesis. Scent dogs should be considered an addition to the gold standard RT-PCR, for rapid testing in situations where great numbers of people from different origins come together. The accuracies may be increased by extending the training phase and selecting individual dogs with better scent detection accuracy. As with any testing scenario, human and in this case dog daily performance could vary. This also applies to the most accurate diagnostic performance of the gold standard RT-PCR that can only be achieved under ideal conditions, which does not always reflects the real life situation. Peer reviewed and preliminary systematic reviews indicate PCR sensitivities ranging from 71 to 100%^29,30^ implying false negative results ranging up to 29% under real-life conditions.

In order to generate rapid test results, a large number of over-the-counter rapid antigen tests are currently used. Test results are generated within about 15 minutes. According to the manufacturers, the tests approved in Germany have diagnostic sensitivities between 91 and 98% and specificities between 98 and 100%^31^. However, the diagnostic accuracy under real-life conditions is estimated to be much lower under these conditions (pre-prints^32,33^). The Paul Ehrlich Institute (Langen, Germany) specified minimum criteria for approved rapid antigen test for SARS-CoV-2 infections. They require a diagnostic sensitivity of above 80% and specificity above 97%^34^. The scent dog method would meet these criteria.

## Conclusions

Detection dogs were able to transfer the conditioned scent detection of BPL-inactivated saliva samples to non-inactivated saliva, urine and sweat samples, with a sensitivity >80% and specifity >94%. All three fluids were equally suited for SARS-CoV-2 detection by dogs and could be used for disease specific VOCs’ pattern recognition. Detection dogs may provide a reliable screening method for SARS-CoV-2 infections in various settings to generate immediate results that can be verified by the gold standard (RT-PCR). Further work, especially under real-life conditions in settings where many individuals have to be screened is needed to fully evaluate the potential of the dog detection method.

## Supporting information

Supplementary Attachments

## List of abbreviations

RT-PCR: reverse transcription polymerase chain reaction test
VOCs: volatile organic compounds
DDTS: Dog Detection Training System
BPL: beta-propiolactone
TP: true positive
FP: false positive
TN: true negative
FN: false negative
CI: confidence interval
TADD: Training Aid Delivery Device

## Declarations

### Ethics approval and consent to participate

The study was conducted according to the ethical requirements established by the Declaration of Helsinki. The local Ethics Committee of *Hannover Medical School* (MHH) and Hamburg Medical Association for the *University Medical-Center Hamburg-Eppendorf* (UKE) approved the study (ethic consent number 9042_BO_K_2020 and PV7298, respectively). Written consent from all participants was collected before sample collection.

## Authors’ contributions

PJ participated in the planning of the study, carried out the main practical work, data analyses and drafted the manuscript. FT, SM and HAV designed and coordinated the study, drafted the manuscript, conducted and coordinated (FT) the sample acquisition and were responsible for data analyses. MvKB and AbO participated in the planning of the laboratory part of the study and were in charge for the legal permission for sample processing. CS and VP carried out the laboratory work including sample preparation, virus inactivation and RT-PCR. JE and HE programmed the DDTS software and HE was also supporting the dog training. IP, TW, MPM, AF and MMA were in charge for the ethical approval, patient recruitment and sample collection (IP, AF) at Hannover Medical School (IP, TW, MPM) and University Medical-Center Hamburg-Eppendorf (AF, MMA). CE was responsible for the special research proposal of the German Armed Forces whereas WS was responsible for the general medical care of the dogs. As project manager on the part of the German Armed Forces, ME coordinated the cooperation with the University of Veterinary Medicine Hannover. ES was responsible for the dog training and helped with data analyses. ME and ES were also involved in designing and coordinating the study. AP, KM and US established and validated the RT-PCR-assays including sampling, sample storage and sample preparation and did the diagnostics. All authors have read and approved the final manuscript.

## Acknowledgements

We would like to thank the members of the ID-UKE-COVID-19 Study Group (Amelie Alberti, Marylyn M. Addo, Etienne Bartels, Thomas T. Brehm, Christine Dahlke, Anahita Fathi, Monika Friedrich, Svenja Hardtke, Till Koch, Ansgar W. Lohse, My L. Ly, Stefan Schmiedel, L. Marie Weskamm, Julian Schulze zur Wiesch) at the University Medical-Center Hamburg-Eppendorf by helping us with recruitment of patients and sample collection. We further would like to thank **S**ina Knisel, Carlos Acosta, Robert Zacharz, Christian Mertel, Pascal Baum, Matthias Wichow, and Jannik Hofmann for support at the University of Veterinary Medicine Hannover, Germany during dog training and Leander Buchner from Bundeswehr Medical Service Headquarters, German Armed Forces, Germany for the support in getting samples. A special thanks goes to Hans Ebbers, Kynoscience, for the support regarding the DDTS. Special thanks go to our “doggy noses” Donnie, Lotta, Coyote, Füge, Vine, Filou, Bellatrix, Margo, Erec Junior and Floki. We would also like to thank Rouwen Stucke and Katharina Meyer for their support in the laboratory and Karin Lübbert for her technical assistance. Heartfelt thanks go to all the people providing us with samples, especially to the SARS-CoV-2 infected persons and their relatives with the sincere intention to contribute to the containment of COVID-19 and to scientific progress. We wish you lots of strength and full recovery during the current pandemic.

