## Supplementary Attachments for "Scent dog identification of SARS-CoV-2 infections, similar across different body fluids"

**Supplementary table 1.** Characteristics of the samples used for the study

| **Sample ID** | **Sex** | **Age (range in years)** | **Sample material** | **SARS-CoV-2 RT-PCR (swab)** | **SARS-CoV-2 RT-PCR (sample material)** | | | **Symptom status** | **Test type** |
| --- | --- | --- | --- | --- | --- | --- | --- | --- | --- |
|  |  |  |  |  | SARS2-IP4-FAM | internal control EGFP 10^9-ML-HEX | result |  |  |
| PS2 | female | 20-30 | saliva | positive | N/A | N/A | N/A | severe* | Training (inactivated saliva) |
| NK-A | male | 40-50 | saliva | negative | N/A | N/A | N/A | none | Training (inactivated saliva) |
| PS7 | male | 60-70 | saliva | positive | N/A | N/A | N/A | severe* | Training (inactivated saliva) |
| NK-M | male | 50-60 | saliva | negative | N/A | N/A | N/A | none | Training (inactivated saliva) |
| PS8 | female | 40-50 | saliva | positive | N/A | N/A | N/A | severe* | Training (inactivated saliva) |
| NK-N | female | 40-50 | saliva | negative | N/A | N/A | N/A | none | Training (inactivated saliva) |
| PS6 | female | 50-60 | saliva | positive | N/A | N/A | N/A | severe* | Training (inactivated saliva) |
| NK-K | female | 50-60 | saliva | negative | N/A | N/A | N/A | none | Training (inactivated saliva) |
| PS9 | male | 60-70 | saliva | positive | N/A | N/A | N/A | severe* | Training (inactivated saliva) |
| NK-E | female | 30-40 | saliva | negative | N/A | N/A | N/A | none | Training (inactivated saliva) |
| PS10 | male | Okt 20 | saliva | positive | N/A | N/A | N/A | mild | Training (inactivated saliva) |
| NK-C | male | 30-40 | saliva | negative | N/A | N/A | N/A | none | Training (inactivated saliva) |
| PS11 | female | 40-50 | saliva | positive | N/A | N/A | N/A | mild | Training (inactivated saliva) |
| NK-H | female | 30-40 | saliva | negative | N/A | N/A | N/A | none | Training (inactivated saliva) |
| PS12 | male | 40-50 | saliva | positive | N/A | N/A | N/A | severe* | Training (inactivated saliva) |
| NK-I | female | 30-40 | saliva | negative | N/A | N/A | N/A | none | Training (inactivated saliva) |
| PS13 | male | Okt 20 | saliva | positive | N/A | N/A | N/A | severe* | Training (inactivated saliva) |
| NK-B | male | 30-40 | saliva | negative | N/A | N/A | N/A | none | Training (inactivated saliva) |
| PS14 | male | 70-80 | saliva | positive | N/A | N/A | N/A | severe* | Training (inactivated saliva) |
| NK-Q | male | 50-60 | saliva | negative | N/A | N/A | N/A | none | Training (inactivated saliva) |
| PS15 | male | 20-30 | saliva | positive | N/A | N/A | N/A | severe* | Training (inactivated saliva) |
| NK-F | female | 40-50 | saliva | negative | N/A | N/A | N/A | none | Training (inactivated saliva) |
| PS16 | male | 40-50 | saliva | positive | N/A | N/A | N/A | severe* | Training (inactivated saliva) |
| NK-G | female | 20-30 | saliva | negative | N/A | N/A | N/A | none | Training (inactivated saliva) |
| PS17 | male | Okt 20 | saliva | positive | 33.33 | 33.70 | positive | mild | Transfer inactive to active saliva |
| NK-T | female | 20-30 | saliva | negative | N/A | N/A | N/A | none | Transfer inactive to active saliva |
| PS71 | female | 20-30 | saliva | positive | 32.65 | 32.24 | positive | mild | Transfer inactive to active saliva |
| NK-U | female | 20-30 | saliva | negative | N/A | N/A | N/A | none | Transfer inactive to active saliva |
| T067 | female | 50-60 | saliva | negative | No Cq | 26.60 | negative | none | Transfer inactive to active saliva |
| PS25 | male | 40-50 | saliva | positive | 37.73 | 30.45 | positive | mild | Transfer inactive to active saliva |
| T079 | male | 40-50 | saliva | negative | No Cq | 26.88 | negative | none | Transfer inactive to active saliva |
| PS73 | male | 80-90 | saliva | positive | No Cq | 32.09 | negative | severe* | Transfer inactive to active saliva |
| T060 | male | 50-60 | saliva | negative | No Cq | 30.12 | negative | none | Transfer inactive to active saliva |
| PS30 | female | 20-30 | saliva | positive | 35.98 | 32.64 | positive | mild | Transfer inactive to active saliva |
| PS96 | female | 20-30 | saliva | negative | No Cq | 31.61 | negative | none | Transfer inactive to active saliva |
| PS61 | female | 50-60 | saliva | positive | No Cq | 33.33 | negative | asymptomatic | Transfer inactive to active saliva |
| PS95 | female | 20-30 | saliva | negative | No Cq | 30.48 | negative | none | Transfer inactive to active saliva |
| PS20 | female | 30-40 | saliva | positive | 26.60 | 31.40 | positive | mild | Transfer to urine and sweat |
| T031 | female | 30-40 | saliva | negative | No Cq | 26.53 | negative | none | Transfer to urine and sweat |
| PS21 | female | 40-50 | saliva | positive | No Cq | 31.49 | negative | mild | Transfer to urine and sweat |
| PS46 | female | 40-50 | saliva | negative | No Cq | 31.78 | negative | none | Transfer to urine and sweat |
| PS23 | male | 20-30 | urine | positive | No Cq | 29.85 | negative | mild | Transfer to urine and sweat |
| PS41 | male | 20-30 | urine | negative | No Cq | 29.63 | negative | none | Transfer to urine and sweat |
| PS63 | female | 20-30 | urine | positive | No Cq | 29.69 | negative | mild | Transfer to urine and sweat |
| PS52 | female | 20-30 | urine | negative | No Cq | 28.94 | negative | none | Transfer to urine and sweat |
| PS29 | female | 20-30 | sweat | positive | No Cq | 29.39 | negative | mild | Transfer to urine and sweat |
| PS42 | female | 20-30 | sweat | negative | No Cq | 29.44 | negative | none | Transfer to urine and sweat |
| PS31 | male | 30-40 | sweat | positive | No Cq | 29.70 | negative | mild | Transfer to urine and sweat |
| PS45 | female | 20-30 | sweat | negative | No Cq | 31.45 | negative | none | Transfer to urine and sweat |
| PS35 | female | 70-80 | sweat | positive | 37.35 | 29.17 | positive | asymptomatic | Transfer to urine and sweat |
| PS57 | female | 20-30 | sweat | negative | No Cq | 29.49 | negative | none | Transfer to urine and sweat |
| PS19 | male | Okt 20 | sweat | positive | No Cq | 31.54 | negative | asyptomatic | pure sweat |
| PS93 | female | 20-30 | sweat | negative | No Cq | 29.33 | negative | none | pure sweat |
| PS26 | female | 40-50 | sweat | positive | No Cq | 29.54 | negative | mild | pure sweat |
| PS53 | female | 20-30 | sweat | negative | No Cq | 31.75 | negative | none | pure sweat |
| PS77 | female | 70-80 | sweat | positive | No Cq | 31.36 | negative | severe* | pure sweat |
| PS82 | female | 30-40 | sweat | negative | No Cq | 31.65 | negative | none | pure sweat |
| PS70 | male | 50-60 | sweat | positive | No Cq | 30.07 | negative | severe* | pure sweat |
| PS84 | female | 30-40 | sweat | negative | No Cq | 31.23 | negative | none | pure sweat |
| PS68 | female | 50-60 | sweat | positive | 37.31 | 29.58 | positive | mild | pure sweat |
| PS81 | female | 20-30 | sweat | negative | No Cq | 29.58 | negative | none | pure sweat |
| PS39 | female | 50-60 | sweat | positive | No Cq | 29.94 | negative | mild | pure sweat |
| PS62 | female | 30-40 | sweat | negative (distractor) | No Cq | 29.90 | negative | mild | pure sweat |
| PS64 | male | 30-40 | sweat | positive | No Cq | 29.67 | negative | mild | pure sweat |
| PS85 | male | 20-30 | sweat | negative | No Cq | 32.04 | negative | none | pure sweat |
| PS65 | female | 30-40 | urine | positive | No Cq | 30.63 | negative | mild | pure urine |
| PS49 | female | 20-30 | urine | negative | No Cq | 34.06 | negative | none | pure urine |
| PS22 | male | Okt 20 | urine | positive | No Cq | 31.10 | negative | mild | pure urine |
| PS87 | male | 20-30 | urine | negative | No Cq | 29.68 | negative | none | pure urine |
| PS79 | female | 50-60 | urine | positive | No Cq | 31.08 | negative | mild | pure urine |
| PS86 | female | 20-30 | urine | negative | No Cq | 29.29 | negative | none | pure urine |
| PS36 | female | Okt 20 | urine | positive | No Cq | 29.39 | negative | mild | pure urine |
| PS94 | female | 20-30 | urine | negative | No Cq | 29.42 | negative | none | pure urine |
| PS27 | male | 40-50 | urine | positive | No Cq | 29.18 | negative | mild | pure urine |
| PS66 | male | 30-40 | urine | negative (distractor) | No Cq | 30.46 | negative | mild | pure urine |
| PS60 | male | 60-70 | urine | positive | No Cq | 33.30 | negative | severe | pure urine |
| PS88 | female | 20-30 | urine | negative | No Cq | 29.41 | negative | none | pure urine |
| PS97 | female | 20-30 | urine | positive | No Cq | 29.55 | negative | mild | pure urine |
| PS89 | female | 20-30 | urine | negative | No Cq | 31.42 | negative | none | pure urine |
| PS80 | female | 20-30 | saliva | positive | 28.40 | 29.47 | positive | mild | pure saliva |
| PS67 | female | 30-40 | saliva | negative (distractor) | No Cq | 30.55 | negative | mild | pure saliva |
| PS69 | female | 70-80 | saliva | positive | 30.68 | 32.74 | positive | severe* | pure saliva |
| T050 | female | 50-60 | saliva | negative | No Cq | 30.65 | negative | none | pure saliva |
| PS32 | male | 50-60 | saliva | positive | No Cq | 31.99 | negative | mild | pure saliva |
| T023 | male | 50-60 | saliva | negative | No Cq | 27.93 | negative | none | pure saliva |
| PS37 | female | Okt 20 | saliva | positive | 28.94 | 31.00 | positive | mild | pure saliva |
| T068 | female | Okt 20 | saliva | negative | No Cq | 27.12 | negative | none | pure saliva |
| PS76 | female | 70-80 | saliva | positive | No Cq | 32.44 | negative | severe* | pure saliva |
| T058 | female | 40-50 | saliva | negative | No Cq | 28.51 | negative | none | pure saliva |
| PS72 | female | 60-70 | saliva | positive | 21.27 | 30.48 | positive | severe* | pure saliva |
| PS18 | female | 30-40 | saliva | negative (distractor) | No Cq | 31.22 | negative | mild | pure saliva |
| PS78 | female | 70-80 | saliva | positive | 33.24 | 30.67 | positive | severe* | pure saliva |
| PS47 | female | 20-30 | saliva | negative | No Cq | 31.97 | negative | none | pure saliva |
| *hospitalised |  |  |  |  |  |  |  |  |  |
| N/A not applicable | |  |  |  |  |  |  |  |  |
| Cq quantification cycle | |  |  |  |  |  |  |  |  |

**Supplementary table 2.**  Characteristics of dogs in the study

| **Name** | **Sex** | **Age (years)** | **Breed** | **Specialty** |
| --- | --- | --- | --- | --- |
| Lotta | Female | 5 | Labrador Retriever | Explosive detection dog |
| Coyote | Male, castrated | 9 | Dutch Shepherd Mix | Explosives detection and protection work |
| Donnie | Male | 3 | Malinois | Explosives detection and protection work |
| Filou | Female | 3 | Malinois | Mine detection dog |
| Füge | Female, castrated | 4 | German Shepherd | No previous training except obedience |
| Vine | Female, castrated | 5 | Malinois | No previous training except obedience |
| Bellatrix | Female | 1 | Labrador Retriever | No previous training except obedience |
| Margo | Female | 1 | Labrador Retriever | No previous training except obedience |
| Floki | Male | 3 | Malinois | No previous training except obedience |
| Erec Junior | Male | 3 | Malinois | Explosives detection and protection work |

**Supplementary table 3.** Results of RT-PCR tests of TADD-membranes and dog noses after testing sessions

| **Sample ID** | **Sample material** | **SARS-CoV-2 RT-PCR (swab)** | **SARS-CoV-2 RT-PCR (outside of the membrane)** | | | **Test type** |
| --- | --- | --- | --- | --- | --- | --- |
|  |  |  | SARS2-IP4-FAM | internal control EGFP 10^9-ML-HEX | result |  |
| NK-T | saliva | negative | No Cq | 31.63 | negative | Transfer inactive to active saliva |
| PS71 | saliva | positive | No Cq | 31.42 | negative | Transfer inactive to active saliva |
| NK-U | saliva | negative | No Cq | 33.89 | negative | Transfer inactive to active saliva |
| T067 | saliva | negative | No Cq | 31.55 | negative | Transfer inactive to active saliva |
| PS25 | saliva | positive | No Cq | 32.78 | negative | Transfer inactive to active saliva |
| T079 | saliva | negative | N/A | N/A | N/A | Transfer inactive to active saliva |
| PS73 | saliva | positive | No Cq | 31.99 | negative | Transfer inactive to active saliva |
| T060 | saliva | negative | N/A | N/A | N/A | Transfer inactive to active saliva |
| PS30 | saliva | positive | No Cq | 31.68 | negative | Transfer inactive to active saliva |
| PS96 | saliva | negative | N/A | N/A | N/A | Transfer inactive to active saliva |
| PS61 | saliva | positive | No Cq | 31.35 | negative | Transfer inactive to active saliva |
| PS95 | saliva | negative | N/A | N/A | N/A | Transfer inactive to active saliva |
| PS20 | saliva | positive | No Cq | 31.24 | negative | Transfer to urine and sweat |
| T031 | saliva | negative | No Cq | 31.13 | negative | Transfer to urine and sweat |
| PS21 | saliva | positive | No Cq | 31.85 | negative | Transfer to urine and sweat |
| PS46 | saliva | negative | No Cq | 31.74 | negative | Transfer to urine and sweat |
| PS23 | urine | positive | No Cq | 31.72 | negative | Transfer to urine and sweat |
| PS41 | urine | negative | No Cq | 31.53 | negative | Transfer to urine and sweat |
| PS63 | urine | positive | No Cq | 31.47 | negative | Transfer to urine and sweat |
| PS52 | urine | negative | N/A | N/A | N/A | Transfer to urine and sweat |
| PS29 | sweat | positive | No Cq | 31.29 | negative | Transfer to urine and sweat |
| PS42 | sweat | negative | N/A | N/A | N/A | Transfer to urine and sweat |
| PS31 | sweat | positive | No Cq | 31.38 | negative | Transfer to urine and sweat |
| PS45 | sweat | negative | N/A | N/A | N/A | Transfer to urine and sweat |
| PS35 | sweat | positive | No Cq | 31.98 | negative | Transfer to urine and sweat |
| PS57 | sweat | negative | N/A | N/A | N/A | Transfer to urine and sweat |
| PS19 | sweat | positive | No Cq | 31.61 | negative | pure sweat |
| PS93 | sweat | negative | No Cq | 31.85 | negative | pure sweat |
| PS26 | sweat | positive | No Cq | 31.87 | negative | pure sweat |
| PS53 | sweat | negative | No Cq | 31.89 | negative | pure sweat |
| PS77 | sweat | positive | No Cq | 32.15 | negative | pure sweat |
| PS82 | sweat | negative | No Cq | 31.82 | negative | pure sweat |
| PS70 | sweat | positive | No Cq | 31.51 | negative | pure sweat |
| PS84 | sweat | negative | N/A | N/A | N/A | pure sweat |
| PS68 | sweat | positive | No Cq | 31.24 | negative | pure sweat |
| PS81 | sweat | negative | N/A | N/A | N/A | pure sweat |
| PS39 | sweat | positive | No Cq | 31.10 | negative | pure sweat |
| PS62 | sweat | negative (distractor) | N/A | N/A | N/A | pure sweat |
| PS64 | sweat | positive | No Cq | 31.40 | negative | pure sweat |
| PS85 | sweat | negative | N/A | N/A | N/A | pure sweat |
| PS65 | urine | positive | No Cq | 33.20 | negative | pure urine |
| PS49 | urine | negative | No Cq | 31.79 | negative | pure urine |
| PS22 | urine | positive | No Cq | 31.86 | negative | pure urine |
| PS87 | urine | negative | No Cq | 32.44 | negative | pure urine |
| PS79 | urine | positive | No Cq | 31.37 | negative | pure urine |
| PS86 | urine | negative | No Cq | 31.79 | negative | pure urine |
| PS36 | urine | positive | No Cq | 31.52 | negative | pure urine |
| PS94 | urine | negative | No Cq | 32.18 | negative | pure urine |
| PS27 | urine | positive | No Cq | 31.26 | negative | pure urine |
| PS66 | urine | negative (distractor) | N/A | N/A | N/A | pure urine |
| PS60 | urine | positive | No Cq | 31.58 | negative | pure urine |
| PS88 | urine | negative | N/A | N/A | N/A | pure urine |
| PS97 | urine | positive | N/A | N/A | N/A | pure urine |
| PS89 | urine | negative | N/A | N/A | N/A | pure urine |
| PS80 | saliva | positive | No Cq | 32.40 | negative | pure saliva |
| PS67 | saliva | negative (distractor) | No Cq | 33.74 | negative | pure saliva |
| PS69 | saliva | positive | No Cq | 35.77 | negative | pure saliva |
| T050 | saliva | negative | No Cq | 34.00 | negative | pure saliva |
| PS32 | saliva | positive | N/A | N/A | N/A | pure saliva |
| T023 | saliva | negative | No Cq | 32.35 | negative | pure saliva |
| PS37 | saliva | positive | No Cq | 34.14 | negative | pure saliva |
| T068 | saliva | negative | N/A | N/A | N/A | pure saliva |
| PS76 | saliva | positive | No Cq | 31.97 | negative | pure saliva |
| T058 | saliva | negative | N/A | N/A | N/A | pure saliva |
| PS72 | saliva | positive | No Cq | 34.10 | negative | pure saliva |
| PS18 | saliva | negative (distractor) | No Cq | 35.32 | negative | pure saliva |
| PS78 | saliva | positive | No Cq | 31.80 | negative | pure saliva |
| PS47 | saliva | negative | N/A | N/A | N/A | pure saliva |

| **Dog** | **Testing day** | **SARS-CoV-2 RT-PCR (nasopharyngeal swab from dogs)** | | |
| --- | --- | --- | --- | --- |
|  |  | SARS2-IP4-FAM | internal control EGFP 10^9-ML-HEX | result |
| 1 | 1 | No Cq | 32.20 | negative |
| 2 | 1 | No Cq | 34.45 | negative |
| 3 | 1 | No Cq | 34.88 | negative |
| 4 | 1 | No Cq | 31.25 | negative |
| 5 | 1 | No Cq | 31.43 | negative |
| 6 | 1 | No Cq | 33.44 | negative |
| 7 | 1 | No Cq | 34.27 | negative |
| 8 | 1 | No Cq | 34.24 | negative |
| 9 | 1 | No Cq | 30.94 | negative |
| 10 | 1 | No Cq | 32.06 | negative |
| 1 | 2 | No Cq | 32.08 | negative |
| 2 | 2 | No Cq | 31.16 | negative |
| 3 | 2 | No Cq | 31.32 | negative |
| 4 | 2 | No Cq | 35.22 | negative |
| 5 | 2 | No Cq | 35.20 | negative |
| 6 | 2 | No Cq | 31.49 | negative |
| 7 | 2 | No Cq | 33.32 | negative |
| 8 | 2 | No Cq | 31.38 | negative |
| 9 | 2 | No Cq | 32.45 | negative |
| 10 | 2 | No Cq | 31.49 | negative |
| 1 | 3 | No Cq | 33.09 | negative |
| 2 | 3 | No Cq | 32.16 | negative |
| 3 | 3 | No Cq | 31.12 | negative |
| 4 | 3 | No Cq | 33.45 | negative |
| 5 | 3 | No Cq | 31.53 | negative |
| 6 | 3 | No Cq | 32.07 | negative |
| 7 | 3 | No Cq | 31.86 | negative |
| 8 | 3 | No Cq | 31.34 | negative |
| 9 | 3 | No Cq | 31.15 | negative |
| 10 | 3 | No Cq | 32.28 | negative |
| 1 | 4 | No Cq | 33.85 | negative |
| 2 | 4 | No Cq | 31.40 | negative |
| 3 | 4 | No Cq | 31.51 | negative |
| 4 | 4 | No Cq | 31.40 | negative |
| 5 | 4 | No Cq | 32.24 | negative |
| 6 | 4 | No Cq | 32.91 | negative |
| 7 | 4 | No Cq | 32.81 | negative |
| 8 | 4 | No Cq | 32.62 | negative |
| 9 | 4 | No Cq | 31.94 | negative |
| 10 | 4 | No Cq | 31.27 | negative |
| negative control | all days | No Cq | 32.09 | negative |
| positive control | all days | 29.29236301 | 40.45 | positive |

**Supplementary table 4.** Diagnostic performance of the scent detection dogs

| **Session** | **Dog** | **Detection** | **SARS-CoV-2 infection status** | | **Total number of sample presentations** | **Diagnostic specificity  (Sp)** | **Diagnostic sensitivity  (Se)** | **Confidence interval (95% CI) Sp** | **Confidence interval (95% CI) Se** | **Positive predictive value (PPV)** | **Negative predictive value (NPV)** | **Confidence interval (95% CI) PPV** | **Confidence interval (95% CI) NPV** | **Accuracy** |  |
| --- | --- | --- | --- | --- | --- | --- | --- | --- | --- | --- | --- | --- | --- | --- | --- |
|  |  |  | **positive** | **negative** |  |  |  |  |  |  |  |  |  |  |  |
| **Non-inactivated saliva samples (after 1 week of training with inactivated saliva samples)** | Dog 1 | Yes | 15 | 5 | 132 | 0.9537 | 0.625 | 0.89618–0.98007 | 0.4271–0.78841 | 0.75 | 0.91964 | 0.5313–0.88814 | 0.8543–0.95715 | 0.89394 |  |
|  |  | No | 9 | 103 |  |  |  |  |  |  |  |  |  |  |  |
|  | Dog 2 | Yes | 14 | 5 | 95 | 0.9375 | 0.93333 | 0.8619–0.97301 | 0.70183–0.99658 | 0.73684 | 0.98684 | 0.51208–0.88194 | 0.92916–0.99933 | 0.93684 |  |
|  |  | No | 1 | 75 |  |  |  |  |  |  |  |  |  |  |  |
|  | Dog 3 | Yes | 16 | 5 | 136 | 0.95614 | 0.72727 | 0.90142–0.98112 | 0.51848–0.86849 | 0.7619 | 0.94783 | 0.54909–0.89372 | 0.89083–0.97587 | 0.91912 |  |
|  |  | No | 6 | 109 |  |  |  |  |  |  |  |  |  |  |  |
|  | Dog 4 | Yes | 9 | 12 | 79 | 0.8209 | 0.75 | 0.71253–0.89446 | 0.46769–0.91106 | 0.42857 | 0.94828 | 0.2447–0.63453 | 0.85861–0.9859 | 0.81013 |  |
|  |  | No | 3 | 55 |  |  |  |  |  |  |  |  |  |  |  |
|  | Dog 5 | Yes | 16 | 3 | 81 | 0.95161 | 0.84211 | 0.86712–0.98681 | 0.62435–0.9448 | 0.84211 | 0.95161 | 0.62435–0.9448 | 0.86712–0.98681 | 0.92593 |  |
|  |  | No | 3 | 59 |  |  |  |  |  |  |  |  |  |  |  |
|  | Dog 6 | Yes | 16 | 5 | 107 | 0.94318 | 0.84211 | 0.8738–0.97549 | 0.62435–0.9448 | 0.7619 | 0.96512 | 0.54909–0.89372 | 0.90239–0.99049 | 0.92523 |  |
|  |  | No | 3 | 83 |  |  |  |  |  |  |  |  |  |  |  |
|  | Dog 7 | Yes | 20 | 0 | 114 | 1 | 0.83333 | 0.95906–1 | 0.64147–0.93321 | 1 | 0.95745 | 0.83887–1 | 0.89564–0.98333 | 0.96491 |  |
|  |  | No | 4 | 90 |  |  |  |  |  |  |  |  |  |  |  |
|  | Dog 8 | Yes | 17 | 3 | 92 | 0.96 | 1 | 0.88887–0.9891 | 0.81568–1 | 0.85 | 1 | 0.63958–0.94763 | 0.94935–1 | 0.96739 |  |
|  |  | No | 0 | 72 |  |  |  |  |  |  |  |  |  |  |  |
|  | Dog 9 | Yes | 17 | 4 | 102 | 0.95238 | 0.94444 | 0.88387–0.98133 | 0.74243–0.99715 | 0.80952 | 0.98765 | 0.59999–0.92332 | 0.93333–0.99937 | 0.95098 |  |
|  |  | No | 1 | 80 |  |  |  |  |  |  |  |  |  |  |  |
|  | Dog 10 | Yes | 14 | 7 | 129 | 0.93396 | 0.6087 | 0.86992–0.96765 | 0.40786–0.77842 | 0.66667 | 0.91667 | 0.45373–0.82805 | 0.84917–0.95554 | 0.87597 |  |
|  |  | No | 9 | 99 |  |  |  |  |  |  |  |  |  |  |  |
|  |  |  |  |  |  | **Median Sp** | **Median Se** | **95% CI of median  Sp** | **95% CI of median  Se** | **Median PPV** | **Median NPV** | **95% CI of median  PPV** | **95% CI of median NPV** | **Median accuracy** | **95% CI of median accuracy** |
|  |  |  |  |  |  | **0.952** | **0.83772** | **0.93396–0.96** | **0.625–0.94444** | **0.7619** | **0.95453** | **0.66667–0.85** | **0.91964–0.98765** | **0.92558** | **0.87597–0.96491** |
| **Session** | **Dog** | **Detection** | **SARS-CoV-2 infection status** | | **Total number of sample presentations** | **Diagnostic specificity  (Sp)** | **Diagnostic sensitivity  (Se)** | **Confidence interval (95% CI) Sp** | **Confidence interval (95% CI) Se** | **Positive predictive value (PPV)** | **Negative predictive value (NPV)** | **Confidence interval (95% CI) PPV** | **Confidence interval (95% CI) NPV** | **Accuracy** |  |
|  |  |  | **positive** | **negative** |  |  |  |  |  |  |  |  |  |  |  |
| **Non-inactivated sweat samples** | Dog 1 | Yes | 10 | 0 | 67 | 1 | 0.55556 | 0.9273–1 | 0.33716–0.7544 | 1 | 0.85965 | 0.72247–1 | 0.74676–0.92713 | 0.8806 |  |
|  |  | No | 8 | 49 |  |  |  |  |  |  |  |  |  |  |  |
|  | Dog 2 | Yes | 10 | 4 | 57 | 0.91489 | 1 | 0.80068–0.96641 | 0.72247–1 | 0.71429 | 1 | 0.45351–0.88279 | 0.91799–1 | 0.92982 |  |
|  |  | No | 0 | 43 |  |  |  |  |  |  |  |  |  |  |  |
|  | Dog 3 | Yes | 10 | 5 | 69 | 0.90909 | 0.71429 | 0.80423–0.96054 | 0.45351–0.88279 | 0.66667 | 0.92593 | 0.41714–0.84824 | 0.82446–0.97082 | 0.86957 |  |
|  |  | No | 4 | 50 |  |  |  |  |  |  |  |  |  |  |  |
|  | Dog 4 | Yes | 10 | 3 | 62 | 0.94231 | 1 | 0.84357–0.98428 | 0.72247–1 | 0.76923 | 1 | 0.49744–0.9182 | 0.9273–1 | 0.95161 |  |
|  |  | No | 0 | 49 |  |  |  |  |  |  |  |  |  |  |  |
|  | Dog 5 | Yes | 10 | 7 | 71 | 0.87719 | 0.71429 | 0.76754–0.93922 | 0.45351–0.88279 | 0.58824 | 0.92593 | 0.36005–0.78389 | 0.82446–0.97082 | 0.84507 |  |
|  |  | No | 4 | 50 |  |  |  |  |  |  |  |  |  |  |  |
|  | Dog 6 | Yes | 10 | 4 | 65 | 0.92157 | 0.71429 | 0.815–0.96908 | 0.45351–0.88279 | 0.71429 | 0.92157 | 0.45351–0.88279 | 0.815–0.96908 | 0.87692 |  |
|  |  | No | 4 | 47 |  |  |  |  |  |  |  |  |  |  |  |
|  | Dog 7 | Yes | 10 | 1 | 44 | 0.9697 | 0.90909 | 0.84681–0.99845 | 0.62264–0.99534 | 0.90909 | 0.9697 | 0.62264–0.99534 | 0.84681–0.99845 | 0.95455 |  |
|  |  | No | 1 | 32 |  |  |  |  |  |  |  |  |  |  |  |
|  | Dog 8 | Yes | 10 | 1 | 56 | 0.97778 | 0.90909 | 0.88433–0.99886 | 0.62264–0.99534 | 0.90909 | 0.97778 | 0.62264–0.99534 | 0.88433–0.99886 | 0.96429 |  |
|  |  | No | 1 | 44 |  |  |  |  |  |  |  |  |  |  |  |
|  | Dog 9 | Yes | 10 | 1 | 40 | 0.96552 | 0.90909 | 0.82824–0.99823 | 0.62264–0.99534 | 0.90909 | 0.96552 | 0.62264–0.99534 | 0.82824–0.99823 | 0.95 |  |
|  |  | No | 1 | 28 |  |  |  |  |  |  |  |  |  |  |  |
|  |  |  |  |  |  | **Median Sp** | **Median Se** | **95% CI of median  Sp** | **95% CI of median  Se** | **Median PPV** | **Median NPV** | **95% CI of median  PPV** | **95% CI of median NPV** | **Median accuracy** | **95% CI of median accuracy** |
|  |  |  |  |  |  | **0.94231** | **0.90909** | **0.90909–0.97778** | **0.71429–1** | **0.76923** | **0.96552** | **0.66667–0.90909** | **0.92157–1** | **0.92982** | **0.86957–0.95455** |
| **Session** | **Dog** | **Detection** | **SARS-CoV-2 infection status** | | **Total number of sample presentations** | **Diagnostic specificity  (Sp)** | **Diagnostic sensitivity  (Se)** | **Confidence interval (95% CI) Sp** | **Confidence interval (95% CI) Se** | **Positive predictive value (PPV)** | **Negative predictive value (NPV)** | **Confidence interval (95% CI) PPV** | **Confidence interval (95% CI) NPV** | **Accuracy** |  |
|  |  |  | **positive** | **negative** |  |  |  |  |  |  |  |  |  |  |  |
| **Non-inactivated urine samples** | Dog 1 | Yes | 10 | 1 | 91 | 0.98592 | 0.5 | 0.92444–0.99928 | 0.2993–0.7007 | 0.90909 | 0.875 | 0.62264–0.99534 | 0.78497–0.93066 | 0.87912 |  |
|  |  | No | 10 | 70 |  |  |  |  |  |  |  |  |  |  |  |
|  | Dog 2 | Yes | 10 | 2 | 50 | 0.94872 | 0.90909 | 0.83114–0.99089 | 0.62264–0.99534 | 0.83333 | 0.97368 | 0.55197–0.97039 | 0.86505–0.99865 | 0.94 |  |
|  |  | No | 1 | 37 |  |  |  |  |  |  |  |  |  |  |  |
|  | Dog 3 | Yes | 10 | 1 | 63 | 0.98 | 0.76923 | 0.89505–0.99897 | 0.49744–0.9182 | 0.90909 | 0.94231 | 0.62264–0.99534 | 0.84357–0.98428 | 0.93651 |  |
|  |  | No | 3 | 49 |  |  |  |  |  |  |  |  |  |  |  |
|  | Dog 4 | Yes | 8 | 1 | 72 | 0.98333 | 0.66667 | 0.91145–0.99915 | 0.39062–0.86188 | 0.88889 | 0.93651 | 0.565–0.9943 | 0.84781–0.97503 | 0.93056 |  |
|  |  | No | 4 | 59 |  |  |  |  |  |  |  |  |  |  |  |
|  | Dog 5 | Yes | 10 | 2 | 49 | 0.94872 | 1 | 0.83114–0.99089 | 0.72247–1 | 0.83333 | 1 | 0.55197–0.97039 | 0.90594–1 | 0.95918 |  |
|  |  | No | 0 | 37 |  |  |  |  |  |  |  |  |  |  |  |
|  | Dog 6 | Yes | 10 | 2 | 66 | 0.96429 | 1 | 0.87881–0.99365 | 0.72247–1 | 0.83333 | 1 | 0.55197–0.97039 | 0.93359–1 | 0.9697 |  |
|  |  | No | 0 | 54 |  |  |  |  |  |  |  |  |  |  |  |
|  | Dog 7 | Yes | 10 | 0 | 41 | 1 | 0.90909 | 0.88649–1 | 0.62264–0.99534 | 1 | 0.96774 | 0.72247–1 | 0.83806–0.99835 | 0.97561 |  |
|  |  | No | 1 | 30 |  |  |  |  |  |  |  |  |  |  |  |
|  | Dog 8 | Yes | 10 | 0 | 47 | 1 | 1 | 0.90594–1 | 0.72247–1 | 1 | 1 | 0.72247–1 | 0.90594–1 | 1 |  |
|  |  | No | 0 | 37 |  |  |  |  |  |  |  |  |  |  |  |
|  | Dog 9 | Yes | 10 | 0 | 49 | 1 | 1 | 0.91033–1 | 0.72247–1 | 1 | 1 | 0.72247–1 | 0.91033–1 | 1 |  |
|  |  | No | 0 | 39 |  |  |  |  |  |  |  |  |  |  |  |
|  | Dog 10 | Yes | 10 | 4 | 66 | 0.92857 | 1 | 0.83025–0.97187 | 0.72247–1 | 0.71429 | 1 | 0.45351–0.88279 | 0.93121–1 | 0.93939 |  |
|  |  | No | 0 | 52 |  |  |  |  |  |  |  |  |  |  |  |
|  |  |  |  |  |  | **Median Sp** | **Median Se** | **95% CI of median  Sp** | **95% CI of median  Se** | **Median PPV** | **Median NPV** | **95% CI of median  PPV** | **95% CI of median NPV** | **Median accuracy** | **95% CI of median accuracy** |
|  |  |  |  |  |  | **0.98167** | **0.95455** | **0.94872–1** | **0.66667–1** | **0.89899** | **0.98684** | **0.83333–1** | **0.93651–1** | **0.94959** | **0.93056–1** |
| **Session** | **Dog** | **Detection** | **SARS-CoV-2 infection status** | | **Total number of sample presentations** | **Diagnostic specificity  (Sp)** | **Diagnostic sensitivity  (Se)** | **Confidence interval (95% CI) Sp** | **Confidence interval (95% CI) Se** | **Positive predictive value (PPV)** | **Negative predictive value (NPV)** | **Confidence interval (95% CI) PPV** | **Confidence interval (95% CI) NPV** | **Accuracy** |  |
|  |  |  | **positive** | **negative** |  |  |  |  |  |  |  |  |  |  |  |
| **Non-inactivated saliva samples** | Dog 1 | Yes | 19 | 1 | 147 | 0.992 | 0.86364 | 0.95608–0.99959 | 0.66665–0.95251 | 0.95 | 0.97638 | 0.76387–0.99744 | 0.93285–0.99356 | 0.97279 |  |
|  |  | No | 3 | 124 |  |  |  |  |  |  |  |  |  |  |  |
|  | Dog 2 | Yes | 20 | 5 | 128 | 0.95098 | 0.76923 | 0.89034–0.97888 | 0.57948–0.88966 | 0.8 | 0.94175 | 0.60869–0.91139 | 0.8787–0.97303 | 0.91406 |  |
|  |  | No | 6 | 97 |  |  |  |  |  |  |  |  |  |  |  |
|  | Dog 3 | Yes | 20 | 7 | 173 | 0.95205 | 0.74074 | 0.90435–0.97658 | 0.55321–0.86830 | 0.74074 | 0.95205 | 0.55321–0.8683 | 0.90435–0.97658 | 0.91908 |  |
|  |  | No | 7 | 139 |  |  |  |  |  |  |  |  |  |  |  |
|  | Dog 4 | Yes | 20 | 4 | 137 | 0.96522 | 0.90909 | 0.91397–0.98639 | 0.72185–0.98385 | 0.83333 | 0.9823 | 0.64147–0.93321 | 0.93776–0.99686 | 0.9562 |  |
|  |  | No | 2 | 111 |  |  |  |  |  |  |  |  |  |  |  |
|  | Dog 5 | Yes | 20 | 5 | 120 | 0.94949 | 0.95238 | 0.88717–0.97824 | 0.77331–0.99756 | 0.8 | 0.98947 | 0.60869–0.91139 | 0.94276–0.99946 | 0.95 |  |
|  |  | No | 1 | 94 |  |  |  |  |  |  |  |  |  |  |  |
|  | Dog 6 | Yes | 20 | 4 | 111 | 0.95506 | 0.90909 | 0.89007–0.98239 | 0.72185–0.98385 | 0.83333 | 0.97701 | 0.64147–0.93321 | 0.92001–0.99592 | 0.94595 |  |
|  |  | No | 2 | 85 |  |  |  |  |  |  |  |  |  |  |  |
|  | Dog 7 | Yes | 18 | 8 | 163 | 0.94074 | 0.64286 | 0.88742–0.96967 | 0.45830–0.79294 | 0.69231 | 0.92701 | 0.50012–0.83499 | 0.87085–0.95987 | 0.88957 |  |
|  |  | No | 10 | 127 |  |  |  |  |  |  |  |  |  |  |  |
|  | Dog 8 | Yes | 20 | 6 | 164 | 0.95556 | 0.68966 | 0.90643–0.97947 | 0.50770–0.82724 | 0.76923 | 0.93478 | 0.57948–0.88966 | 0.8807–0.96531 | 0.90854 |  |
|  |  | No | 9 | 129 |  |  |  |  |  |  |  |  |  |  |  |
|  | Dog 9 | Yes | 20 | 1 | 112 | 0.98901 | 0.95238 | 0.94035–0.99944 | 0.77331–0.99756 | 0.95238 | 0.98901 | 0.77331–0.99756 | 0.94035–0.99944 | 0.98214 |  |
|  |  | No | 1 | 90 |  |  |  |  |  |  |  |  |  |  |  |
|  | Dog 10 | Yes | 19 | 5 | 191 | 0.96875 | 0.6129 | 0.92894–0.98658 | 0.43824–0.76267 | 0.79167 | 0.92814 | 0.5953–0.90755 | 0.87861–0.95842 | 0.91099 |  |
|  |  | No | 12 | 155 |  |  |  |  |  |  |  |  |  |  |  |
|  |  |  |  |  |  | **Median Sp** | **Median Se** | **95% CI of median  Sp** | **95% CI of median  Se** | **Median PPV** | **Median NPV** | **95% CI of median  PPV** | **95% CI of median NPV** | **Median accuracy** | **95% CI of median accuracy** |
|  |  |  |  |  |  | **0.95531** | **0.81644** | **0.94949–0.98901** | **0.64286–0.95238** | **0.8** | **0.96422** | **0.74074–0.9** | **0.92814–0.98901** | **0.93252** | **0.90854–0.97279** |

**Supplementary table 5.** Detection performance and success rates per session and dog

| **Session** | **Dog** | **Detection** | **SARS-CoV-2 infection status** | | | **Total number of right decisions** | **Total number of sample presentations** | **Success rate per dog** |
| --- | --- | --- | --- | --- | --- | --- | --- | --- |
|  |  |  | **positive** | **negative** | |  |  |  |
| **Non-inactivated saliva samples (after 1 week of training with inactivated saliva samples)** | Dog 1 | Yes | 15 | 5 | | 118 | 132 | 89% |
|  |  | No | 9 | 103 | |  |  |  |
|  | Dog 2 | Yes | 14 | 5 | | 89 | 95 | 94% |
|  |  | No | 1 | 75 | |  |  |  |
|  | Dog 3 | Yes | 16 | 5 | | 125 | 136 | 92% |
|  |  | No | 6 | 109 | |  |  |  |
|  | Dog 4 | Yes | 9 | 12 | | 64 | 79 | 81% |
|  |  | No | 3 | 55 | |  |  |  |
|  | Dog 5 | Yes | 16 | 3 | | 75 | 81 | 93% |
|  |  | No | 3 | 59 | |  |  |  |
|  | Dog 6 | Yes | 16 | 5 | | 99 | 107 | 93% |
|  |  | No | 3 | 83 | |  |  |  |
|  | Dog 7 | Yes | 20 | 0 | | 110 | 114 | 96% |
|  |  | No | 4 | 90 | |  |  |  |
|  | Dog 8 | Yes | 17 | 3 | | 89 | 92 | 97% |
|  |  | No | 0 | 72 | |  |  |  |
|  | Dog 9 | Yes | 17 | 4 | | 97 | 102 | 95% |
|  |  | No | 1 | 80 | |  |  |  |
|  | Dog 10 | Yes | 14 | 7 | | 113 | 129 | 88% |
|  |  | No | 9 | 99 | |  |  |  |
|  | All dogs | Yes | 154 | 49 | | 979 | 1067 | 92% |
|  |  | No | 39 | 825 | |  |  |  |
| **Session** | **Dog** | **Detection** | **SARS-CoV-2 infection status** | | | **Total number of right decisions** | **Total number of sample presentations** | **Success rate per dog** |
|  |  |  | **positive** | **negative** | |  |  |  |
| **Non-inactivated saliva, urine and sweat samples** | Dog 1 | Yes | 18 | 7 | | 174 | 191 | 91% |
|  |  | No | 10 | 156 | |  |  |  |
|  | Dog 2 | Yes | 18 | 19 | | 161 | 186 | 87% |
|  |  | No | 6 | 143 | |  |  |  |
|  | Dog 3 | Yes | 20 | 6 | | 135 | 150 | 90% |
|  |  | No | 9 | 115 | |  |  |  |
|  | Dog 4 | Yes | 18 | 12 | | 169 | 197 | 86% |
|  |  | No | 16 | 151 | |  |  |  |
|  | Dog 5 | Yes | 17 | 15 | | 164 | 195 | 84% |
|  |  | No | 16 | 147 | |  |  |  |
|  | Dog 6 | Yes | 19 | 6 | | 117 | 127 | 92% |
|  |  | No | 4 | 98 | |  |  |  |
|  | Dog 7 | Yes | 19 | 1 | | 201 | 223 | 90% |
|  |  | No | 21 | 182 | |  |  |  |
|  | Dog 8 | Yes | 20 | 1 | | 112 | 115 | 97% |
|  |  | No | 2 | 92 | |  |  |  |
|  | Dog 9 | Yes | 18 | 5 | | 136 | 147 | 93% |
|  |  | No | 6 | 118 | |  |  |  |
|  | Dog 10 | Yes | 18 | 8 | | 124 | 139 | 89% |
|  |  | No | 7 | 106 | |  |  |  |
|  | All dogs | Yes | 185 | 80 | | 1493 | 1670 | 89% |
|  |  | No | 97 | 1308 | |  |  |  |
| **Session** | **Dog** | **Detection** | **SARS-CoV-2 infection status** | | | **Total number of right decisions** | **Total number of sample presentations** | **Success rate per dog** |
|  |  |  | **positive** | **negative** | |  |  |  |
| **Non-inactivated sweat samples** | Dog 1 | Yes | 10 | 0 | | 59 | 67 | 88% |
|  |  | No | 8 | 49 | |  |  |  |
|  | Dog 2 | Yes | 10 | 4 | | 53 | 57 | 93% |
|  |  | No | 0 | 43 | |  |  |  |
|  | Dog 3 | Yes | 10 | 5 | | 60 | 69 | 87% |
|  |  | No | 4 | 50 | |  |  |  |
|  | Dog 4 | Yes | 10 | 3 | | 59 | 62 | 95% |
|  |  | No | 0 | 49 | |  |  |  |
|  | Dog 5 | Yes | 10 | 7 | | 60 | 71 | 85% |
|  |  | No | 4 | 50 | |  |  |  |
|  | Dog 6 | Yes | 10 | 4 | | 57 | 65 | 88% |
|  |  | No | 4 | 47 | |  |  |  |
|  | Dog 7 | Yes | 10 | 1 | | 42 | 44 | 95% |
|  |  | No | 1 | 32 | |  |  |  |
|  | Dog 8 | Yes | 10 | 1 | | 54 | 56 | 96% |
|  |  | No | 1 | 44 | |  |  |  |
|  | Dog 9 | Yes | 10 | 1 | | 38 | 40 | 95% |
|  |  | No | 1 | 28 | |  |  |  |
|  | All dogs | Yes | 90 | 26 | | 482 | 531 | 91% |
|  |  | No | 23 | 392 | |  |  |  |
| **Session** | **Dog** | **Detection** | **SARS-CoV-2 infection status** | | | **Total number of right decisions** | **Total number of sample presentations** | **Success rate per dog** |
|  |  |  | **positive** | | **negative** |  |  |  |
| **Non-inactivated urine samples** | Dog 1 | Yes | 10 | 1 | | 80 | 91 | 88% |
|  |  | No | 10 | 70 | |  |  |  |
|  | Dog 2 | Yes | 10 | 2 | | 47 | 50 | 94% |
|  |  | No | 1 | 37 | |  |  |  |
|  | Dog 3 | Yes | 10 | 1 | | 59 | 63 | 94% |
|  |  | No | 3 | 49 | |  |  |  |
|  | Dog 4 | Yes | 8 | 1 | | 67 | 72 | 93% |
|  |  | No | 4 | 59 | |  |  |  |
|  | Dog 5 | Yes | 10 | 2 | | 47 | 49 | 96% |
|  |  | No | 0 | 37 | |  |  |  |
|  | Dog 6 | Yes | 10 | 2 | | 64 | 66 | 97% |
|  |  | No | 0 | 54 | |  |  |  |
|  | Dog 7 | Yes | 10 | 0 | | 40 | 41 | 98% |
|  |  | No | 1 | 30 | |  |  |  |
|  | Dog 8 | Yes | 10 | 0 | | 47 | 47 | 100% |
|  |  | No | 0 | 37 | |  |  |  |
|  | Dog 9 | Yes | 10 | 0 | | 49 | 49 | 100% |
|  |  | No | 0 | 39 | |  |  |  |
|  | Dog 10 | Yes | 10 | 4 | | 62 | 66 | 94% |
|  |  | No | 0 | 52 | |  |  |  |
|  | All dogs | Yes | 98 | 13 | | 562 | 594 | 95% |
|  |  | No | 19 | 464 | |  |  |  |
| **Session** | **Dog** | **Detection** | **SARS-CoV-2 infection status** | | | **Total number of right decisions** | **Total number of sample presentations** | **Success rate per dog** |
|  |  |  | **positive** | **negative** | |  |  |  |
| **Non-inactivated saliva samples** | Dog 1 | Yes | 19 | 1 | | 143 | 147 | 97% |
|  |  | No | 3 | 124 | |  |  |  |
|  | Dog 2 | Yes | 20 | 5 | | 117 | 128 | 91% |
|  |  | No | 6 | 97 | |  |  |  |
|  | Dog 3 | Yes | 20 | 7 | | 159 | 173 | 92% |
|  |  | No | 7 | 139 | |  |  |  |
|  | Dog 4 | Yes | 20 | 4 | | 131 | 137 | 96% |
|  |  | No | 2 | 111 | |  |  |  |
|  | Dog 5 | Yes | 20 | 5 | | 114 | 120 | 95% |
|  |  | No | 1 | 94 | |  |  |  |
|  | Dog 6 | Yes | 20 | 4 | | 105 | 111 | 95% |
|  |  | No | 2 | 85 | |  |  |  |
|  | Dog 7 | Yes | 18 | 8 | | 145 | 163 | 89% |
|  |  | No | 10 | 127 | |  |  |  |
|  | Dog 8 | Yes | 20 | 6 | | 149 | 164 | 91% |
|  |  | No | 9 | 129 | |  |  |  |
|  | Dog 9 | Yes | 20 | 1 | | 110 | 112 | 98% |
|  |  | No | 1 | 90 | |  |  |  |
|  | Dog 10 | Yes | 19 | 5 | | 174 | 191 | 91% |
|  |  | No | 12 | 155 | |  |  |  |
|  | All dogs | Yes | 196 | 46 | | 1347 | 1446 | 93% |
|  |  | No | 53 | 1151 | |  |  |  |
| **Session** | **Dog** | **Detection** | **SARS-CoV-2 infection status** | | | **Total number of right decisions** | **Total number of sample presentations** | **Success rate per dog** |
|  |  |  | **positive** | **negative** | |  |  |  |
| **All sessions** | Dog 1 | Yes | 72 | 14 | | 574 | 628 | 91% |
|  |  | No | 40 | 502 | |  |  |  |
|  | Dog 2 | Yes | 72 | 35 | | 467 | 516 | 91% |
|  |  | No | 14 | 395 | |  |  |  |
|  | Dog 3 | Yes | 76 | 24 | | 538 | 591 | 91% |
|  |  | No | 29 | 462 | |  |  |  |
|  | Dog 4 | Yes | 65 | 32 | | 490 | 547 | 90% |
|  |  | No | 25 | 425 | |  |  |  |
|  | Dog 5 | Yes | 73 | 32 | | 460 | 516 | 89% |
|  |  | No | 24 | 387 | |  |  |  |
|  | Dog 6 | Yes | 75 | 21 | | 442 | 476 | 93% |
|  |  | No | 13 | 367 | |  |  |  |
|  | Dog 7 | Yes | 77 | 10 | | 538 | 585 | 92% |
|  |  | No | 37 | 461 | |  |  |  |
|  | Dog 8 | Yes | 77 | 11 | | 451 | 474 | 95% |
|  |  | No | 12 | 374 | |  |  |  |
|  | Dog 9 | Yes | 75 | 11 | | 430 | 450 | 96% |
|  |  | No | 9 | 355 | |  |  |  |
|  | Dog 10 | Yes | 61 | 24 | | 473 | 525 | 90% |
|  |  | No | 28 | 412 | |  |  |  |
|  | All dogs | Yes | 723 | 214 | | 4863 | 5308 | 92% |
|  |  | No | 231 | 4140 | |  |  |  |
